# Stability of steady-state visual evoked potential contrast response functions

**DOI:** 10.1101/2022.06.08.495412

**Authors:** Ryan T. Ash, Kerry Nix, Anthony M. Norcia

## Abstract

A repeated measure of neural activity that is stable over time when unperturbed is needed to be able to meaningfully measure neuroplastic changes in the brain. With sensory-evoked potentials in particular, repeated presentation of stimuli can generate neuroplasticity by itself under certain conditions. We assessed the repeated-measure within-day and across-day stability of the steady-state visual-evoked potential (ssVEP), a high signal-to-noise electrophysiological readout of neural activity in human visual cortex, in preparation for studies of visual cortical neuroplasticity. Steady-state VEP contrast-sweep responses were measured daily for 4 days (four 20-trial blocks per day, 22 participants). Response amplitudes were stable in individual participants, with measured across-block and across-day coefficients of variation (CV= SD / Mean) of 12±1% and 19±2%, respectively. No consistent changes in response amplitude were observed either across blocks or across days. We conclude that contrast-sweep steady-state VEPs provide a stable human neurophysiological measure well-suited for repeated-measures studies.

## INTRODUCTION

In order to meaningfully measure neuroplastic changes in the brains of individual participants, one must start with a neurophysiological measure that is at least somewhat stable without perturbation. Stable measures of neural activity are difficult to achieve in human participants. Some commonly-used methods to measure neuroplastic changes in humans include motor-evoked potentials with transcranial magnetic stimulation (TMS), BOLD responses with functional MRI, and event-related potentials (ERPs) with electroencephalography (EEG). A commonly used quantifications of repeated-measure stability is the coefficient of variation (C.V., the standard deviation divided by the mean across measurements expressed as a percentage) (Bennett and Miller, 2010). TMS-evoked MEP response amplitudes show a CV of 20-70% (Rösler et al., 2008; Jung et al., 2010; Ammann et al., 2020). Sensory-evoked fMRI BOLD response amplitudes demonstrate a CV of 20-70% (Leontiev and Buxton, 2007; Bennett and Miller, 2010; Malmqvist et al., 2016; Birman and Gardner, 2018; Trevino et al., 2018) within-day. ERPs show an across-session CV of 10-20 under ideal conditions (Goldberg et al., 2002; Willeford et al., 2013). Few studies have even attempted to assess the across-day stability of these modalities.

A less-used method that may have unique utility to measure human neuroplasticity is the steady-state visual-evoked potential (ssVEP), recognized for its high signal-to-noise ratio (Norcia et al., 2015). Estimates of visual acuity from sweep ssVEPs may have a CV as low as 8% (Lauritzen et al., 2004), but the within-participant stability of ssVEP response amplitudes has not been thoroughly assessed to our knowledge.

Achieving stable repeated measures with sensory-evoked potentials is further complicated by the fact that at least under certain conditions repeated presentation of visual or auditory stimuli can induce neuroplasticity that resembles long-term potentiation(LTP) (Sanders et al., 2018; Sumner et al., 2020). Although previous findings are somewhat conflicting (Dias et al., n.d.; Valstad et al., 2020), several labs have demonstrated that showing 9 Hz flash stimuli for 2 minutes and 2 Hz contrast-reversing stimuli for 10 minutes can both produce an LTP-like increase in the amplitude of the visual evoked potential. Whether repeated contrast-sweep stimuli cause an LTP-like effect has not been tested to our knowledge. Additionally, very few in-human studies have measured responses to repeated visual stimulus presentations across multiple consecutive days, even though this has been shown in animals to induce cumulative LTP-like effects (Cooke and Bear, 2010).

Here we quantified the within-day and across-day stability of ssVEP contrast-response functions, and performed a power analysis to estimate the number of participants needed to detect different degrees of neuroplastic changes.

## METHODS

### Data sets

We used a large contrast-sweep ssVEP data set previously generated in our lab (Dmochowski et al., 2015). Ninety trials of contrast-sweep ssVEPs were recorded daily for four days in a row in 22 participants, using high-density 128-channel EEG.

### EEG recording

The details of data collection are available in (Dmochowski et al., 2015). In brief, male and female neurotypical adult participants were recruited from the Stanford neuroscience community.

Participants were asked to wash their hair with shampoo (without conditioner) in the morning before recording. Vision testing was performed to confirm normal or corrected-to-normal vision. The 128-channel EGI HydroCel net saline headcap (sized to the participant’s head) was soaked in KCl solution for 5-10 minutes and placed on the participant’s head. Electrodes were apposed to the scalp and seeded with additional solution to achieve low (<5 kOhm) impedance in >95% of electrodes. Contrast-sweep stimuli, generated with custom PowerDIVA software, were presented in 4 blocks of 20 trials while the participant performed an orthogonal fixation task, for a total of 100-120 trials. Each trial consisted of a square contrast-reversing sinusoidal luminance grating presented to a single hemifield (left or right, randomized for each participant). The contrast-reversal had a square-wave temporal profile and a frequency of 9 Hz (generating a frequency-doubled response at 18 Hz and higher even harmonics). One in ten trials was randomly presented in the contralateral visual field (data not shown here). The stimuli subtended 0 to 10 degrees eccentricity horizontally and -5 to 5 degrees vertically.

Stimuli started at 2% contrast and increased in log steps up to 80% contrast over ten 1-second bins (10 seconds total). There was a 3-4 second intertrial interval. Participants were allowed to take breaks between blocks as needed. Impedance checks occurred 1-2 times over the course of the experiment session. The entire session lasted between 35 and 45 minutes depending on the duration of breaks. EEG was sampled natively at 500 Hz then resampled at 360 Hz to match the monitor refresh rate (72 Hz, 5 data samples per video frame).

### Data analysis

Preprocessing was performed as in previous studies from the lab (Kaestner et al., 2022). Briefly, raw EEG data sets were preprocessed with a 0.3-50 Hz bandpass filter using EGI software. Then our PowerDIVA-XDiva custom signal processing software performed artifact rejection in two steps. First, the continuous filtered data were evaluated according to a sample-by-sample thresholding procedure to locate consistently noisy sensors. These channels were replaced by the average of their six nearest spatial neighbors. The EEG reference was then re-referenced from Cz to the common average of all sensors. Second, 1-second EEG epochs that contained a large percentage of data samples exceeding threshold (30-80 uV) were excluded on a sensor-by-sensor basis.

The ssVEP amplitude was calculated at the 1^st^, 2^nd^, and 4^th^ harmonic by a recursive least squares adaptive filter (Tang and Norcia, 1995). The filter consisted of two weights, one for the imaginary and one for the real coefficient of each harmonic frequency. Weights were adjusted to minimize the squared estimation error between the reference and recorded signal. The memory length of the filter was 1 second, such that the learned coefficients were averaged over an exponential forgetting function that was equivalent to the duration of one bin of the disparity sweep.

Reliable Components Analysis (RCA) was used for dimensionality reduction (Dmochowski et al., 2015). This technique optimizes the weighting of individual electrodes to maximize trial to trial consistency of the phase-locked ssVEPs. Components were learned on RLS-filtered complex value data from 2F1 and 4F1 responses measured across all trials and all participants. The first Reliable Component (RC1) localized to the contralateral occipital electrodes representing early visual cortex as expected, and this RC was used for all subsequent analysis (Results highly similar to what is reported here were obtained when RCA components were calculated for individual participants). Response amplitude was calculated on each trial by taking the square root of the sum of the squared real and squared imaginary components of the response.

The repeated-measures within-participant coefficient of variation (CV) was calculated as follows: For the within-day/across-block condition, responses were first averaged within block (20 trials). Then, the S.D. was taken across the 4 per-block mean values, divided by the mean across all 4 blocks, and multiplied by 100. For the across-day condition, mean responses were calculated per day (average across 90 trials). Then the S.D. was taken across the 4 per-day mean values, divided by the mean across all 4 days, and multiplied by 100.

## RESULTS

In the ssVEP paradigm, a visual stimulus is presented at a specific flicker frequency (**Fig. 1A**). EEG recorded from visual cortex synchronizes at that frequency, such that a Fourier transform can filter out neural activity unrelated to the visual stimulus (**Fig. 1B**, adapted from Ji et al 2019 for illustration purposes). **Fig. 1C** shows the sensor weights for the right-hemifield stimulus calculated by RCA. Robust responses linearly related to the log of the visual stimulus contrast are observable even in single 10-second contrast-sweep trials (**Fig. 1D**).

**Figure 1.**
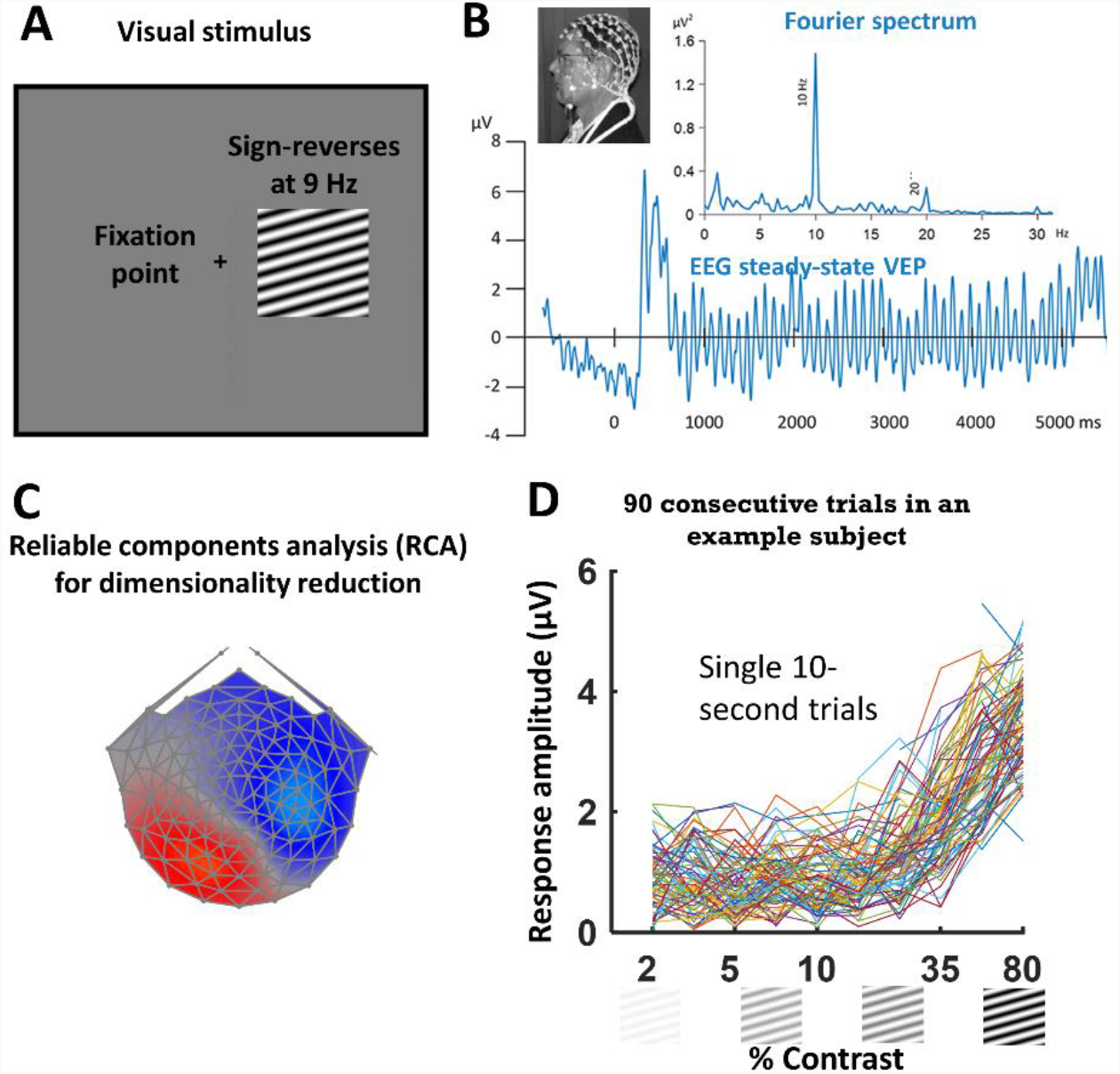
Recording steady state visual-evoked potentials. **A**. Schematic of the visual stimulus presentation. The participant maintains fixation at the fixation point while a contrast-reversing luminance grating (10°, 3 cycles/deg grating stimulus, 9-Hz contrast-reversal) is presented to the left or right visual hemifield in a 10-second sweep from 5 to 80% contrast in 10 log-spaced bins. The hemifield is randomized for each participant and maintained across 4 days of recording. **B**. EEG signals recorded from occipital electrodes oscillate at the frequency of the stimulus contrast alteration rate. A Fourier Transform allows the EEG response to the stimulus to be quantified (inset). This plot adapted from (Ji et al., 2019) for illustration purposes only. **C**. Reliable components analysis (RCA) is used to identify the electrodes whose signals phase-lock to the stimulus and to aggregate these electrodes for dimensionality reduction (see Methods). This image visualizes a top-view of the electrode topography, where red heatmap color identifies the electrodes that synchronize to the right hemifield stimulus, in the contralateral occipital area as expected. **D**. Response amplitude by contrast for 90 10-second trial responses from an example session and single participant, showing that responses are robust on the single trial level and vary similarly with contrast across trials.

We first assessed the within-day stability across 80 trials (split into 4 20-trial blocks, with the first 10 of 90 trials excluded as the participant settled into the fixation task), recorded over ∼30 minutes. High-SNR contrast response functions were observable in most participants, and these were quite similar across the 4 blocks of recording (**Fig. 2A**, black and gray lines), with a mild decrease in the response over time observable in some examples. When contrast-response functions were plotted across 4 days of recording, some changes in response were seen from day to day, but the overall shapes of the contrast-response function were similar across days in individuals (**Fig. 2B**).

**Figure 2.**
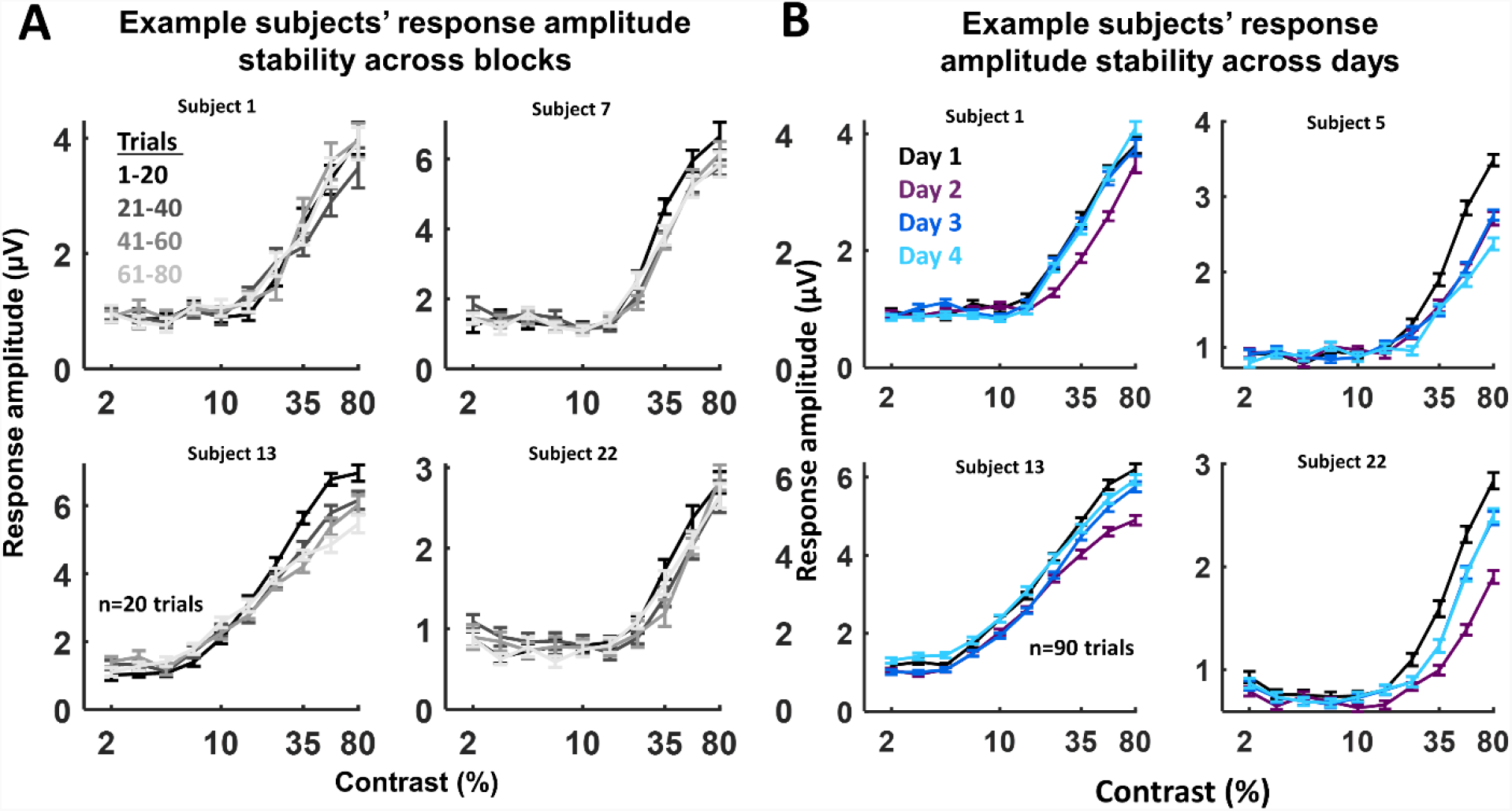
Single-participant contrast-response functions across blocks and days. **A**. Contrast response functions (CRFs) plotted across four 20-trial blocks for four example participants (each panel). Grayscale color depicts block number. Contrast response functions are highly stable across the ∼45 minute recording session. With slight inconsistent changes of the response over time. **B**. CRFs plotted across 4 consecutive days of recording for four example participants. Line color depicts recording day (see legend). Contrast-response function shape is strongly maintained across days, with some fluctuation in response amplitude across days.

As a shorthand for plotting the response amplitude across participants, we determined the response at the peak of the contrast-response function (Rmax), defined as the activity at the average response at the two adjacent contrasts with the highest mean response, and at the middle of the contrast response function (Rmid), defined as the average response of the two contrasts where the response was closest to half of the max response (see Methods). Average (thick black line, error bars S.D.) and per-participant (gray lines) Rmid and Rmax responses were visualized across the four 20-trial blocks (**Fig. 3A**) and four days (**Fig. 3B**) of recording. This visualization demonstrates that for most participants, the responses were similar across the four blocks or days of recording, with a subset of participants showing decreases or increases inconsistently across recording sessions.

**Figure 3.**
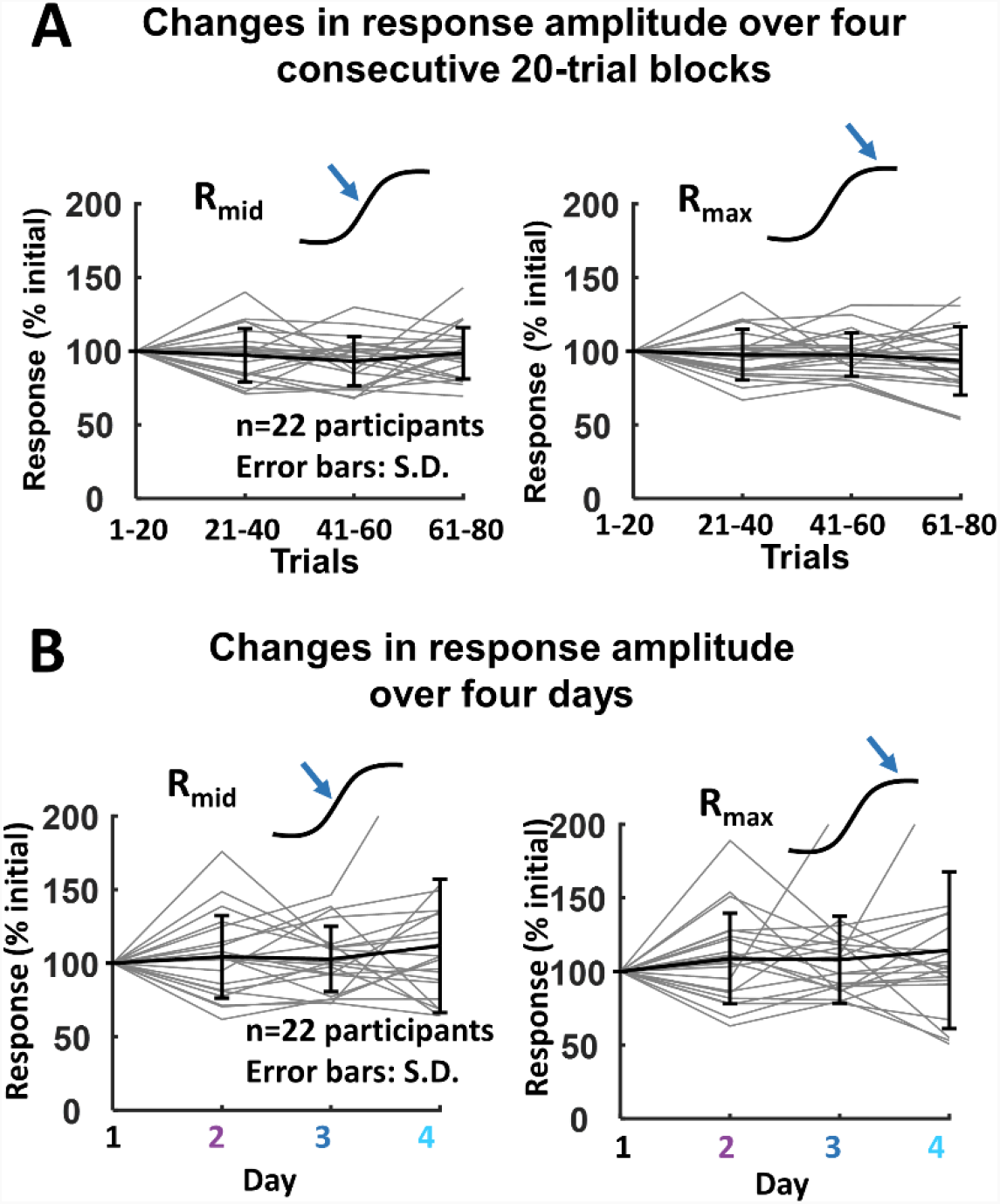
Summary data of changes in response amplitude across blocks and days. **A**. Per-block response amplitudes were normalized to the initial block (Trials 1-20) and plotted over time at the middle (left panel) and max of the contrast response function (right panel). The stimulus contrasts that represented Rmid and Rmax were individualized for each participant. Thin gray lines show single-participant data, and thick lines show mean +/-SD across participants (n=22). **B**. Day-1 normalized response amplitudes per day at the middle and max of the CRF, plotted as in A, across 4 days of recording.

To quantify the stability of the response amplitude over time, we calculated the repeated-measure coefficient of variation (CV), which is the standard deviation of the response across blocks or days normalized by the mean response across blocks/days, multiplied by 100 and expressed as a percentage. The across-block and across-day CV were 12±1 and 19±2 respectively at Rmid, 12±1 and 20±2, respectively at Rmax (**Fig. 4A**), indicating that ssVEP response amplitude is generally stable to within 12% across blocks and 20% across days.

**Figure 4.**
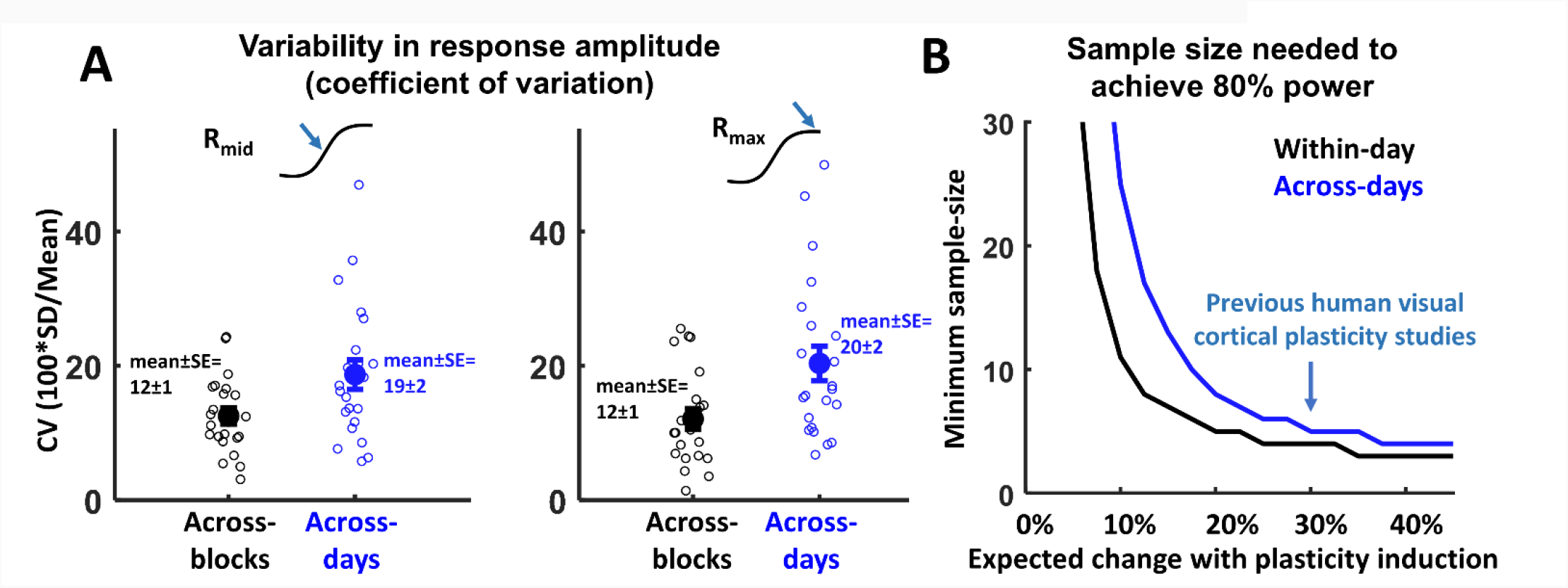
ssVEP response amplitude coefficient of variation and power analysis. **A**. The coefficient of variation (100* S.D./Mean) of the response amplitude across blocks (black) and across days (blue) was plotted per participant (small circles) and as mean +/-SE across participants (large circles), for Rmax. Similar values were observed for Rmid (data not shown). **B**. Visualization of the sample size needed to detect neuroplastic changes based on the measured CV across blocks (black) and across days (blue) at 80% statistical power. The plot shows that neuroplastic changes of 30% (as seen in previous studies including (Bohotin et al., 2002; Kirk et al., 2010) may be detectable with as few as 5-7 participants given the measured response stability.

A power analysis was performed to assess the sample size needed to detect a significant change in ssVEP response amplitude such as following a plasticity protocol, given our measured response variability. **Fig. 4B** shows the minimum sample-size needed to achieve a statistical power of 0.8 as a function of the expected response amplitude change with plasticity induction. Previous in vivo measurements of human visual cortical neuroplasticity generally show a change in activity post-plasticity-induction of 20-40% (Thut and Pascual-Leone, 2010; Kirk et al., 2020). Our statistical power calculation estimates that the contrast-sweep ssVEP is powered to measure changes in response amplitude with as few as 6-7 participants (**Fig. 4B, teal arrow**), providing strong evidence that if there were a neuroplastic change in the current experiment we would have been well powered to detect it.

## DISCUSSION

Here we report that response amplitudes measured with repeated presentations of the contrast-sweep ssVEP are stable over time, across trials within-day and across days. Response amplitudes vary with a coefficient of variation of 12% across 4 blocks within a ∼45 min recording session, and 19-20% across 4 days of consecutive recording. The signals were stable over multiple sources of variability including variation of EEG electrode positioning over days, fluctuations in participant brain state factors including arousal and attention, variability in level of task engagement and fixation (although at a gross level task performance was monitored during experiment), neuronal sensitization and synaptic plasticity in response to repeated sensory input, and other drifts in thalamocortical excitability. These factors can change both within and across recording session, and all have been shown to affect neural activity and EEG responses (Luck, 2014; Norcia et al., 2015). Nonetheless, we observed remarkably stable responses.

Our measured CVs of 12% across blocks within day and 19-20% across days suggest that ssVEPs have superior stability compared to some commonly used human neurophysiological measures, including MEPs, which show an across-block CV of 20-70% (Rösler et al., 2008; Jung et al., 2010; Ammann et al., 2020) and sensory-evoked fMRI BOLD responses which also show an across-block CV of 20-70% (Leontiev and Buxton, 2007; Bennett and Miller, 2010; Malmqvist et al., 2016; Birman and Gardner, 2018; Trevino et al., 2018). Our CV measurements compare favorably to that measured ERPs, which under ideal conditions show an across-block CV of 10-20 (Goldberg et al., 2002; Willeford et al., 2013), and EEG TMS-evoked potentials, which show an across-block CV of 15-20 (Van Doren et al., 2015), see also (Kerwin et al., 2018). Together the data suggest that ssVEP response amplitudes are among the more stable human neurophysiological measures currently available to neuroscientists and clinicians.

Although others have shown a paradoxical increase in ERP response amplitude following repetitive “tetanic” sensory stimulation reflecting LTP-like processes (Cooke and Bear, 2010; Sanders et al., 2018; Sumner et al., 2020), we did not observe this pattern in our data. We expect that this may be due to differences in details of the visual stimulus and EEG read-out we used. First, the temporal frequency of the inducing stimulus may be a key factor, with prior paradigms using a stimulus modulation rate of 2 Hz or ∼9 Hz; our stimuli had a reversal rate of 18 Hz. A somewhat simplified summary of previous work in this space suggests that 2 minutes of flashed checkerboards/gratings at 9 Hz most reliably induced LTP, while 2, 5, and 18 Hz were less effective (Normann et al., 2007; Kirk et al., 2020). It has been theorized that 9 Hz induces LTP most effectively because it is close to the resonant frequency of visual cortex (i.e. alpha) (Sumner et al., 2020). For contrast-reversing stimuli, 2 Hz but not 19 Hz was able to induce sensory LTP (Normann et al., 2007; Sumner et al., 2020). Apparently, for flashed-onset stimuli the higher frequencies are more effective at inducing sensory LTP, while for contrast-reversal stimuli the lower frequencies are more effective. Presumably this is related to how the different stimuli synchronously activate volleys of synapses in the visual cortex. The stimulus used in the current study was contrast-reversing at 18 Hz. Therefore it was faster than stimuli previously shown to generate LTP, particularly as is was contrast-reversing instead of flash-onset stimulus, which may only be expected to induce LTP-like effects at 2 Hz (Normann et al., 2007). Second, most previous studies have read out effects of tetanic stimulation with transient ERPs rather than the ssVEP to the inducing stimulus as we do here. Finally, the other major difference is that our inducing stimulus comprised a 10-second contrast-sweep with 3-5 second inter trial interval, vs. either 2 or 10 minutes of continuous repetitive stimulation in the other sensory LTP protocols.

Additionally, it is important to note along these lines that there is large variability in the sensory LTP literature. Some have questioned if sensory LTP is even a valid construct (Dias et al., 2022), and the largest studies have shown modest if any consistent effects across hundreds of participants (Valstad et al., 2020). Our finding of response stability with repeated stimulus presentation provides further support for VEP stability, but a more thorough investigation of stimulus parameters is needed to determine in what regime ssVEP stimuli do or do not generate neuroplastic effects.

The stability of the ssVEP creates a novel opportunity for within-participant studies of neuroplasticity in relatively small numbers of participants, including repetitive sensory neuroplasticity processes (Cooke and Bear, 2010; Kirk et al., 2010) and repetitive brain stimulation-induced neuroplasticity (Thut and Pascual-Leone, 2010; Darmani et al., 2022).

